# Cross-reactive neutralization of SARS-CoV-2 by serum antibodies from recovered SARS patients and immunized animals

**DOI:** 10.1101/2020.04.20.052126

**Authors:** Yuanmei Zhu, Danwei Yu, Yang Han, Hongxia Yan, Huihui Chong, Lili Ren, Jianwei Wang, Taisheng Li, Yuxian He

**Author notes:** Correspondence to J.W., T. L., or Y.H. These authors contributed equally to this work (Yuanmei Zhu, Danwei Yu, Yang Han).

## Abstract

The current COVID-19 pandemic, caused by a novel coronavirus SARS-CoV-2, poses serious threats to public health and social stability, calling for urgent need for vaccines and therapeutics. SARS-CoV-2 is genetically close to SARS-CoV, thus it is important to define the between antigenic cross-reactivity and neutralization. In this study, we firstly analyzed 20 convalescent serum samples collected from SARS-CoV infected individuals during the 2003 SARS outbreak. All patient sera reacted strongly with the S1 subunit and receptor-binding domain (RBD) of SARS-CoV, cross-reacted with the S ectodomain, S1, RBD, and S2 proteins of SARS-CoV-2, and neutralized both SARS-CoV and SARS-CoV-2 S protein-driven infections. Multiple panels of antisera from mice and rabbits immunized with a full-length S and RBD immunogens of SARS-CoV were also characterized, verifying the cross-reactive neutralization against SARS-CoV-2. Interestingly, we found that a palm civet SARS-CoV-derived RBD elicited more potent cross-neutralizing responses in immunized animals than the RBD from a human SARS-CoV strain, informing a strategy to develop a universe vaccine against emerging CoVs.

**Summary:** Serum antibodies from SARS-CoV infected patients and immunized animals cross-neutralize SARS-CoV-2 suggests strategies for universe vaccines against emerging CoVs.

## Introduction

The global outbreak of the Coronavirus Disease 2019 (COVID-19) was caused by severe acute respiratory syndrome coronavirus 2 (SARS-CoV-2), which is a new coronavirus (CoV) genetically close to SARS-CoV emerged in 2002 (*1-3*). As of 20 April 2020, a total of 2,246,291 confirmed COVID-19 cases, including 152,707 deaths, have been reported from 213 countries or regions, and the numbers are growing rapidly (https://www.who.int). The pandemic threatens to become one of the most difficult times faced by humans in modern history. Unfortunately, even though 17 years passed, we have not developed effective prophylactics and therapeutics in preparedness for the re-emergence of SARS or SARS-like CoVs. A vaccine is urgently needed to prevent the human-to-human transmission of SARS-CoV-2.

Like SARS-CoV and many other CoVs, SARS-CoV-2 utilizes its surface spike (S) glycoprotein to gain entry into host cells (*4-6*). Typically, the S protein forms a homotrimer with each protomer consisting of S1 and S2 subunits. The N-terminal S1 subunit is responsible for virus binding to the cellular receptor ACE2 through an internal receptor-binding domain (RBD) that is capable of functional folding independently, whereas the membrane-proximal S2 subunit mediates membrane fusion events. Very recently, the prefusion structure of the SARS-CoV-2 S protein was determined by cryo-EM, which revealed an overall similarity to that of SARS-CoV (*4, 7*); the crystal structure of the SARS-CoV-2 RBD in complex with ACE2 was also determined by several independent groups, and the residues or motifs critical for the higher-affinity RBD-ACE2 interaction were identified (*8-10*).

The S protein of CoVs is also a main target of neutralizing antibodies (nAbs) thus being considered an immunogen for vaccine development (*4, 11*). During the SARS-CoV outbreak in 2002, we took immediate actions to characterize the immune responses in infected SARS patients and in inactivated virus vaccine- or S protein-immunized animals (*12-20*). We demonstrated that the S protein RBD dominates the nAb response against SARS-CoV infection and thus proposed a RBD-based vaccine strategy (*11, 15-22*). Our follow-up studies verified a potent and persistent anti-RBD response in recovered SARS patients (*23-25*). Although SARS-CoV-2 and SARS-CoV share substantial genetic and functional similarities, their S proteins, especially in the RBD sequences, display relatively larger divergences. Toward developing vaccines and immunotheraptics against emerging CoVs, it is fundamentally important to characterize the antigenic cross-reactivity between SARS-CoV-2 and SARS-CoV.

## Results

### Serum antibodies from recovered SARS patients react strongly with the S protein of SARS-CoV-2

A panel of serum samples collected from 20 patients recovered from SARS-CoV infection was analyzed for the antigenic cross-reactivity with SARS-CoV-2. Firstly, we examined the convalescent sera by a commercial diagnostic ELISA kit, which uses a recombinant nucleocapsid (N) protein of SARS-CoV-2 as detection antigen. As shown in Figure 1A, all the serum samples at a 1:100 dilution displayed high reactivity, verifying that the N antigen is highly conserved between SARS-CoV and SARS-CoV-2. As tested by ELISA, each of the patient sera also reacted with the SARS-CoV S1 subunit and its RBD strongly (Fig. 1B). Then, we determined the cross-reactivity of the patient sera with four recombinant protein antigens derived from the S protein of SARS-CoV-2, including S ectodomain (designated S), S1 subunit, RBD, and S2 subunit. As shown in Fig. 1C, all the serum samples also reacted strongly with the S and S2 proteins, but they were less reactive with the S1 and RBD proteins.

**Figure 1.**
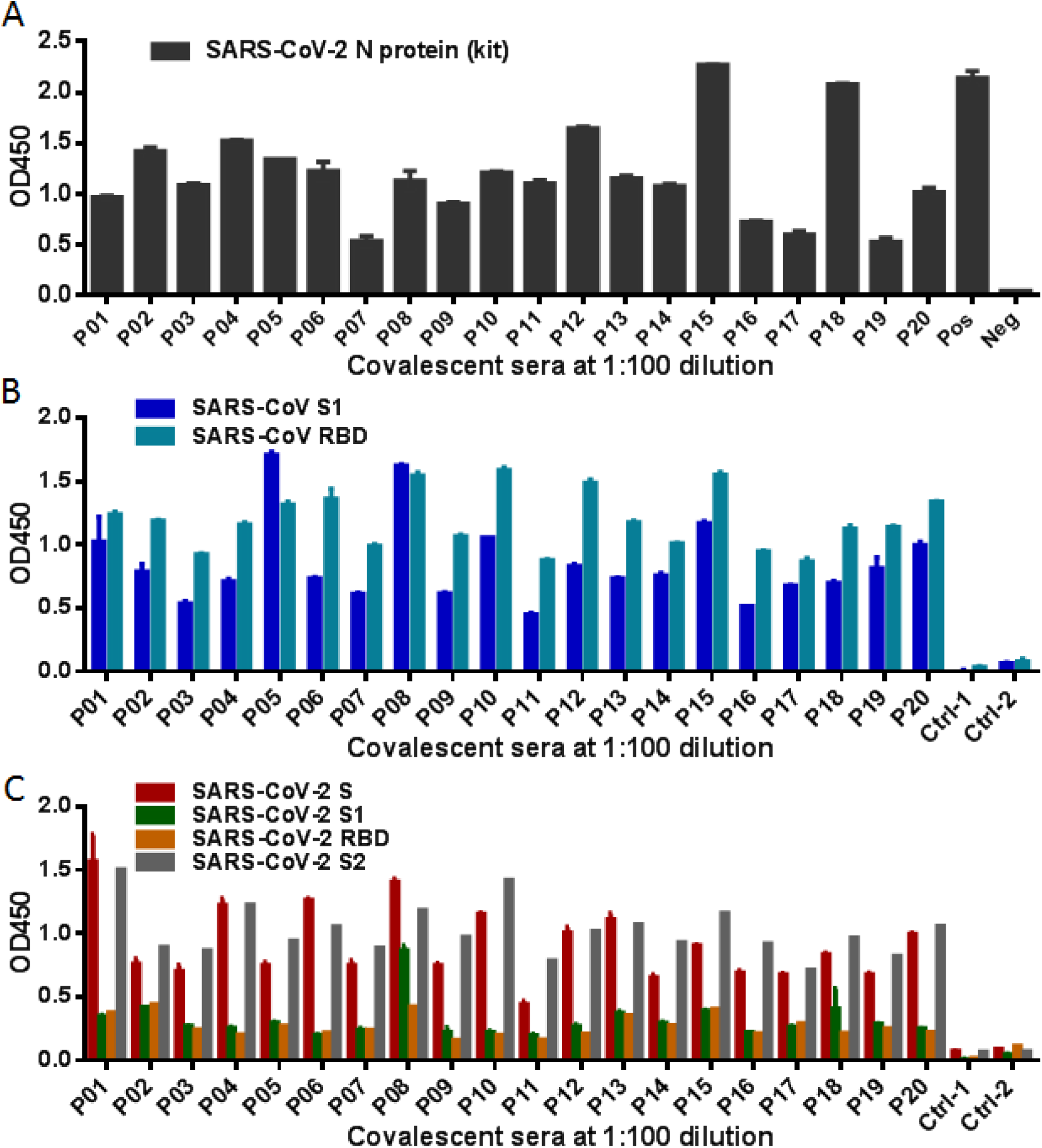
Cross-reactivity of convalescent sera from SARS-CoV infected patients with SARS-CoV-2 determined by ELISA. **(A)** Reactivity of sera from 20 recovered SARS-CoV patients (P01 to P20) with the nucleoprotein (N) of SARS-CoV-2 was measured by a commercial ELISA kit. **(B)** Reactivity of convalescent SARS sera with the recombinant S1 and RBD proteins of SARS-CoV. **(C)** Reactivity of convalescent SARS sera with the S ectodomain (designated S), S1, RBD, and S2 proteins of SARS-CoV-2. Serum samples from two healthy donors were used as negative control (Ctrl-1 and Ctrl-2). The experiments were performed with duplicate samples and repeated three times, and data are shown as means with standard deviations.

### Serum antibodies from recovered SARS patients cross-neutralize SARS-CoV-2

As limited by facility that can handle authentic viruses, we developed a pseudovirus-based single-cycle infection assay to determine the cross-neutralizing activity of the convalescent SARS sera on SARS-CoV and SARS-CoV-2. The pseudotype for vesicular stomatitis virus (VSV-G) was also prepared as a virus control. Initially, the serum samples were analyzed at a 1:20 dilution. As shown in Fig. 2A, all the sera efficiently neutralized both the SARS-CoV and SARS-CoV-2 pseudoviruses to infect 239T/ACE2 cells, and in comparison, each serum had lower efficiency in inhibiting SARS-CoV-2 as compared to SARS-CoV. None of the immune sera showed appreciable neutralizing activity on VSV-G pseudovirus. The neutralizing titer for each patient serum was then determined. As shown in Fig. 2B, the patient sera could neutralize SARS-CoV with titers ranging from 1:120 to 1: 3,240 and cross-neutralized SARS-CoV-2 with titers ranging from 1:20 to 1: 360. In highlight, the patient P08 serum had the highest titer to neutralize SARS-CoV (1: 3,240) when it neutralized SARS-CoV-2 with a titer of 1:120; the patient P13 serum showed the highest titer on SARS-CoV-2 (1:360) when it had a 1:1,080 titer to efficiently neutralize SARS-CoV.

**Figure 2.**
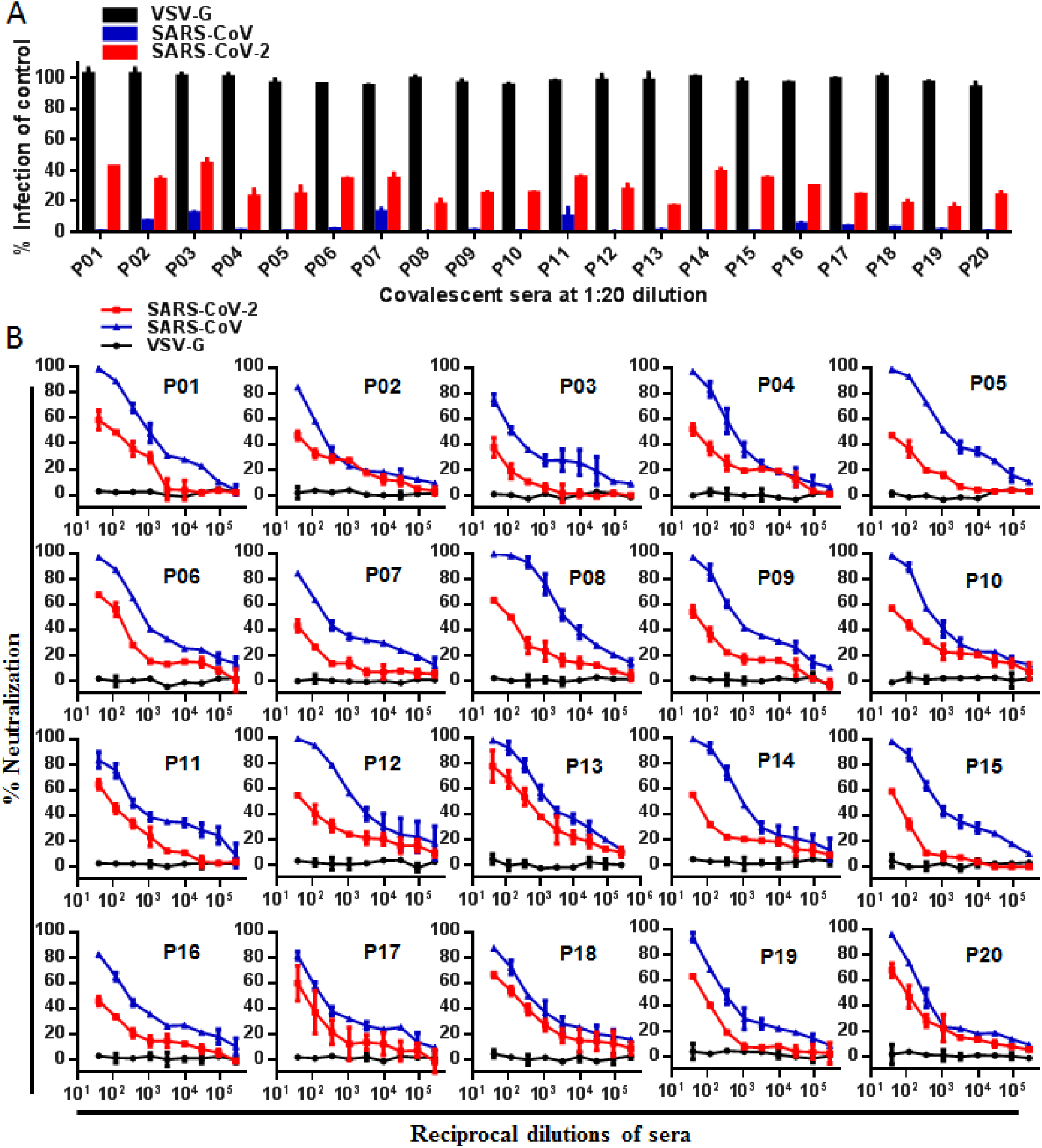
Neutralizing activity of convalescent sera from SARS patients against SARS-CoV and SARS-CoV-2 determined by single-cycle infection assay. **(A)** Neutralizing activity of convalescent patient sera was tested at a 1:20 dilution. Statistical significance was tested by two-way ANOVA with Dunnett posttest, indicating that all the sera significantly inhibited SARS-CoV and SARS-CoV-2 (p<0.001) but not VSV-G (p>0.05) pseudoviruses to infect 293T/ACE2 cells. **(B)** Neutralizing titers of each of convalescent patient sera on three pseudotypes were measured. The experiments were performed with triplicate samples and repeated three times, and data are shown as means with standard deviations.

### Mouse antisera raised against SARS-CoV S protein react and neutralize SARS-CoV-2

To comprehensively characterize the cross-reactivity between the S proteins of SARS-CoV and SARS-CoV-2, we generated mouse antisera against the S protein of SARS-CoV by immunization. Herein, three mice (M-1, M-2, and M-3) were immunized with a recombinant full-length S protein in the presence of MLP-TDM adjuvant, while two mice (M-4 and M-5) were immunized with the S protein plus alum adjuvant. Binding of antisera to diverse S antigens were initially examined by ELISA. As shown in Fig. 3A, the mice immunized by the S protein with the MLP-TDM adjuvant developed relatively higher titers of antibody responses as compared to the two mice with the alum adjuvant. Apparently, each of mouse antisera had high cross-reactivity with the SARS-CoV-2 S and S2 proteins, but the cross-reactive antibodies specific for the SARS-CoV-2 S1 and RBD were hardly detected except the anti-S1 response in mouse M3. Subsequently, the neutralizing capacity of mouse anti-S sera was measured with pseudoviruses. As shown in Fig. 3B to 3F, all the antisera, diluted at 1: 40, 1: 160, or 1: 640, potently neutralized SARS-CoV, and consistently, they were able to cross-neutralize SARS-CoV-2 although with reduced capacity relative to SARS-CoV.

**Figure 3.**
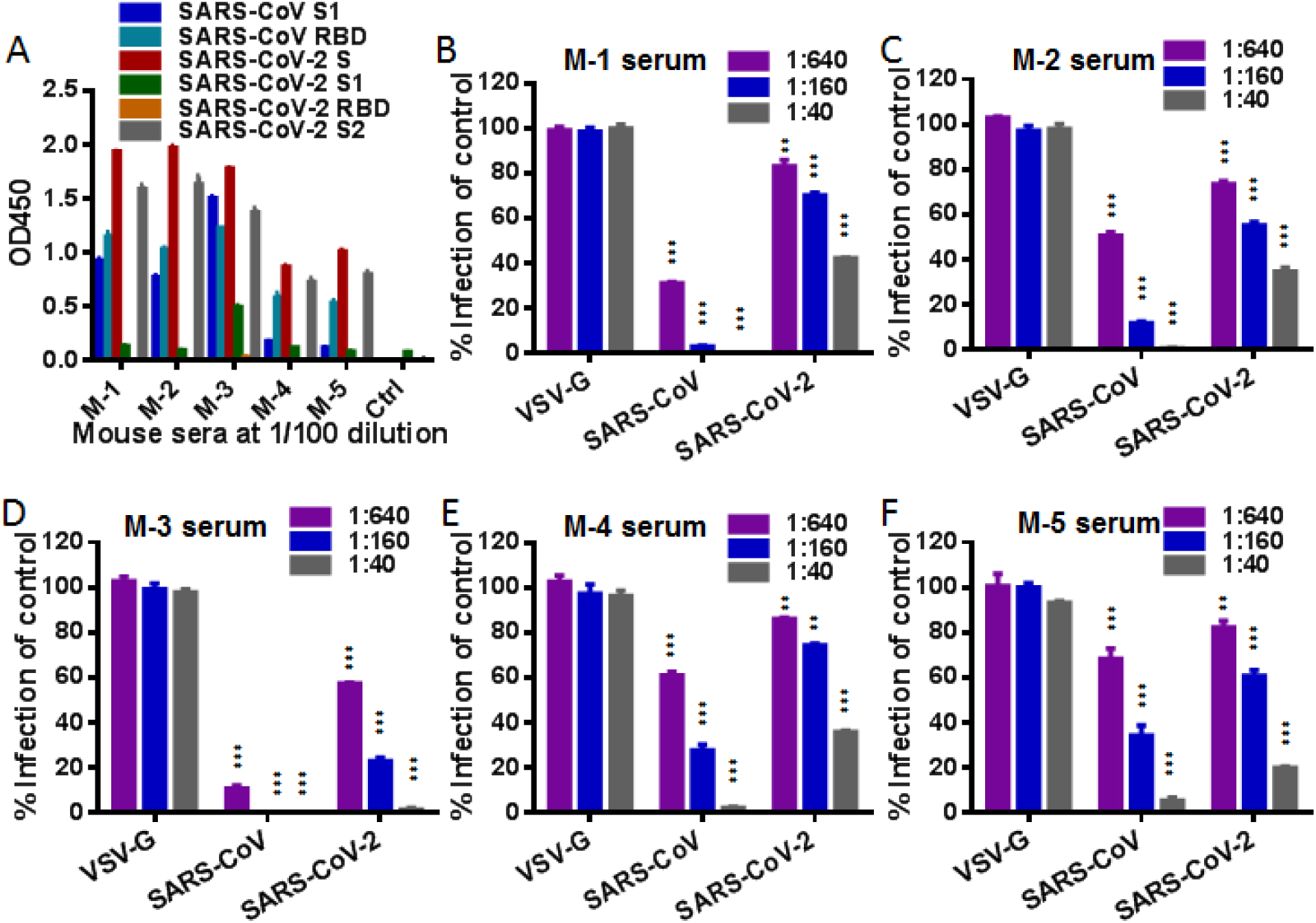
Cross-reactive and neutralizing activities of antisera from mice immunized with a full-length S protein of SARS-CoV. **(A)** Binding activity of mouse antisera at a 1:100 dilution to SARS-CoV (S1 and RBD) and SARS-CoV-2 (S, S1, RBD, and S2) antigens was determined by ELISA. A healthy mouse serum was tested as control. **(B)** Neutralizing activity of mouse antisera at indicated dilutions against SARS-CoV, SARS-CoV-2, and VSV-G pseudoviruses was determined by a single-cycle infection assay. The experiments were performed in triplicates and repeated three times, and data are shown as means with standard deviations. Statistical significance was tested by two-way ANOVA with Dunnett posttest.

### Mouse and rabbit antisera developed against SARS-CoV RBD cross-react and neutralize SARS-CoV-2

As the S protein RBD dominates the nAb response to SARS-CoV, we sought to characterize the RBD-mediated cross-reactivity and neutralization on SARS-CoV-2. To this end, we firstly generated mouse anti-RBD sera by immunization with two RBD-Fc fusion proteins: one encoding the RBD sequence of a palm civet SARS-CoV strain SZ16 (SZ16-RBD) and the second one with the RBD sequence of a human SARS-CoV strain GD03 (GD03-RBD). As shown in Fig. 4A, all of eight mice developed robust antibody responses against the SARS-CoV S1 and RBD; and in comparison, four mice (m1 to m4) immunized with SZ16-RBD exhibited higher titers of antibody responses than the mice (m5 to m8) immunized with GD03-RBD. Each of anti-RBD sera cross-reacted well with the S protein of SARS-CoV-2, suggesting that SARS-CoV and SARS-CoV-2 do share antigenically conserved epitopes in the RBD sites. Noticeably, while the SZ16-RBD immune sera also reacted with the SARS-CoV-2 RBD antigen, the reactivity of the GD03-RBD immune sera was hardly detected, implying that recombinant RBD protein of SARS-CoV-2 used here might not be correctly folded to mimic the antigenic conformation presented on its S protein. Similarly, the neutralizing activity of mouse antisera was determined by pseudovirus-based single-cycle infection assay. As shown by Fig. 4B to 4I, each antiserum even at a high dilution (1:640) displayed very potent activity to neutralize SARS-CoV; they also cross-neutralized SARS-CoV-2 with relatively lower efficiency.

**Figure 4.**
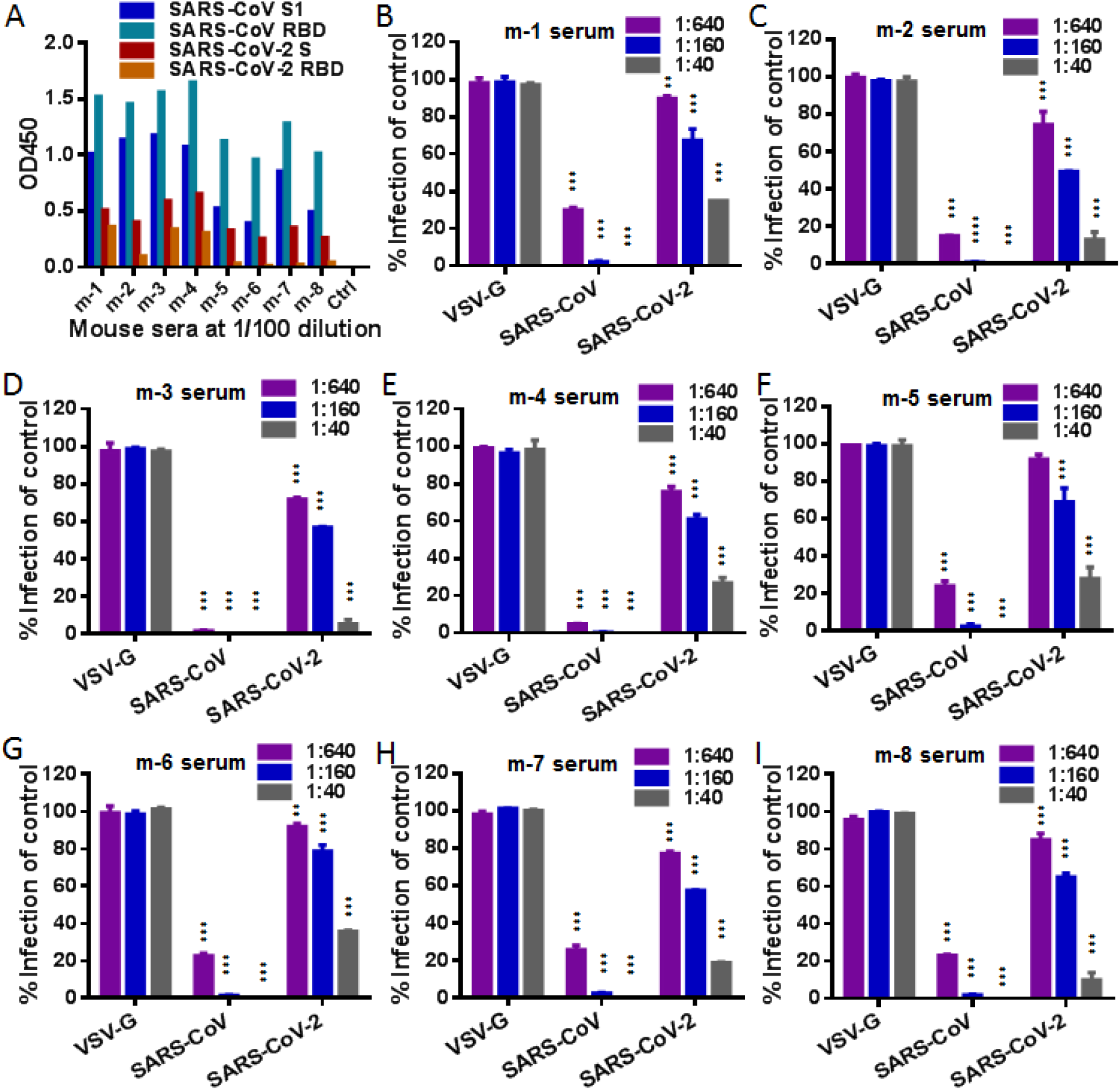
Cross-reactive and neutralizing activities of antisera from mice immunized with RBD proteins of SARS-CoV. **(A)** Binding activity of mouse antisera at a 1:100 dilution to SARS-CoV (S1 and RBD) and SARS-CoV-2 (S protein and RBD) antigens was determined by ELISA. A healthy mouse serum was tested as control. **(B)** Neutralizing activity of mouse antisera at indicated dilutions against SARS-CoV, SARS-CoV-2, and VSV-G pseudoviruses was determined by a single-cycle infection assay. The experiments were performed in triplicates and repeated three times, and data are shown as means with standard deviations. Statistical significance was tested by two-way ANOVA with Dunnett posttest.

We further developed rabbit antisera by immunizations, in which two rabbits were immunized with SZ16-RBD and two rabbits were with GD03-RBD. Interestingly, both RBD proteins elicited antibodies highly reactive with both the SARS-CoV and SARS-CoV-2 antigens (Fig. S1), which were different from their immunizations in mice. As expected, all of the rabbit antisera potently neutralized both SARS-CoV and SARS-CoV-2 in a similar profile with that of the mouse anti-S and anti-RBD sera. Again, the results verified that the SARS-CoV S protein and its RBD immunogens can induce cross-neutralizing antibodies toward SARS-CoV-2 by vaccination.

### Rabbit antibodies induced by SZ16-RBD but not GD03 can block RBD binding to 293T/ACE2 cells

To validate the observed cross-reactive neutralization and explore the underlying mechanism, we purified anti-RBD antibodies from the rabbit antisera above. As shown in Fig. 5A, four purified rabbit anti-RBD antibodies reacted strongly with the SARS-CoV RBD protein and cross-reacted with the SARS-CoV-2 S and RBD but not S2 proteins in a dose-dependent manner. Consistent to its antiserum, the purified rabbit R-4 antibody was less reactive with the SARS-CoV-2 antigens. Moreover, the purified antibodies dose-dependently neutralized SARS-CoV and SARS-CoV-2 but not VSV-G (Fig. 5B). Next, we investigated whether the rabbit anti-RBD antibodies block RBD binding to 293T/ACE2 cells by flow cytometry. As expected, both the SARS-CoV and SARS-CoV-2 RBD proteins could bind to 293T/ACE2 cells in a dose-dependent manner, and in a line with previous findings that the RBD of SARS-CoV-2 bind to ACE2 more efficiently (Fig. S2). Surprisingly, the antibodies purified from SZ16-RBD-immunized rabbits (R-1 and R-2) potently blocked the binding of both the RBD proteins, whereas the antibodies from GD03-RBD-immunized rabbits (R-3 and R-4) had no such blocking functionality except a high concentration of the rabbit R-3 antibody on the SARS-CoV RBD binding (Fig. 6).

**Figure 5.**
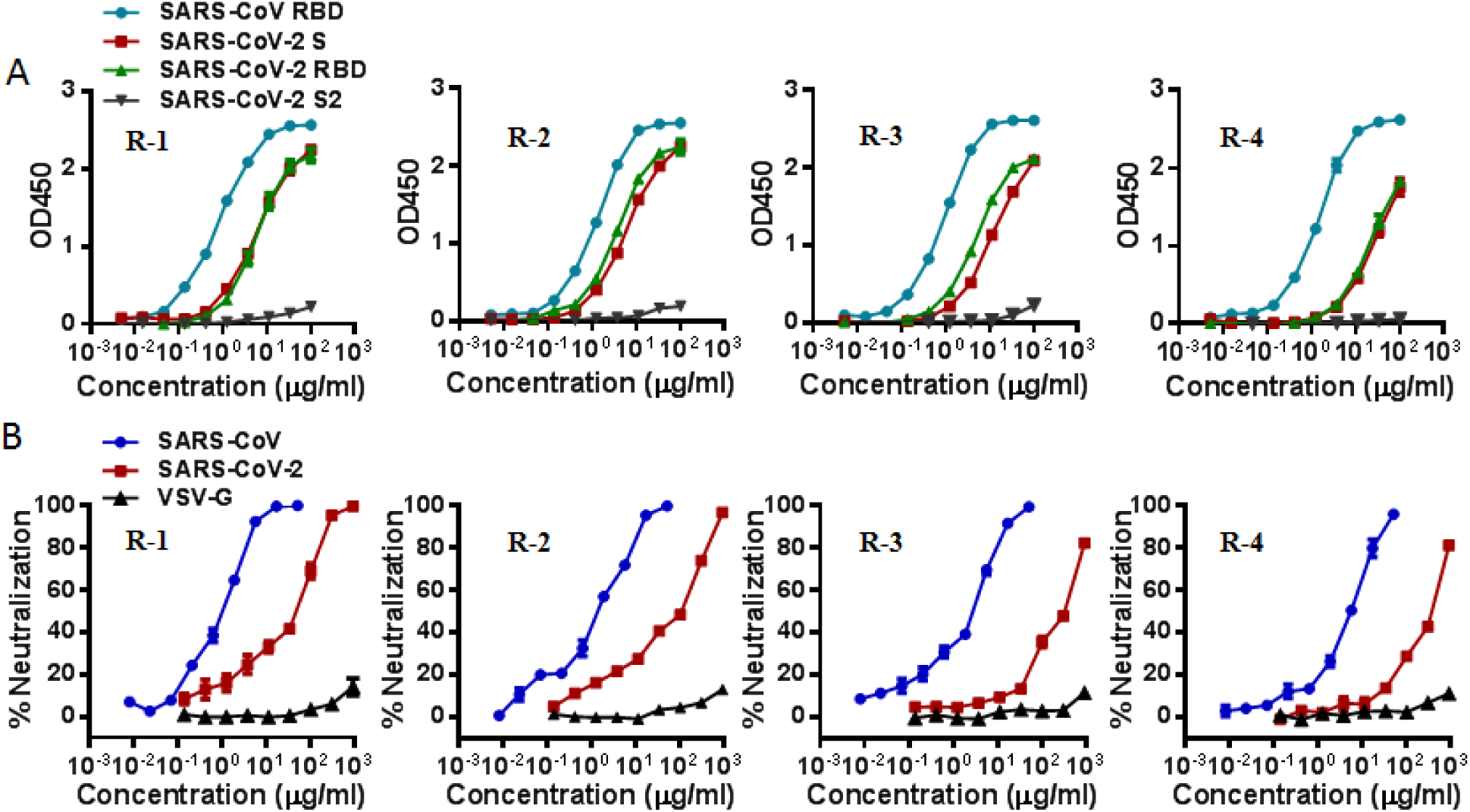
Cross-reactivity and neutralization of purified rabbit anti-RBD antibodies. **(A)** Binding titers of purified rabbit anti-RBD antibodies to SARS-CoV (RBD) and SARS-CoV-2 (S, RBD, and S2) antigens were determined by ELISA. A healthy rabbit serum was tested as control. **(B)** Neutralizing titers of purified rabbit anti-RBD antibodies on SARS-CoV, SARS-CoV-2, and VSV-G pseudoviruses was determined by a single-cycle infection assay. The experiments were done in triplicates and repeated three times, and data are shown as means with standard deviations.

**Figure 6.**
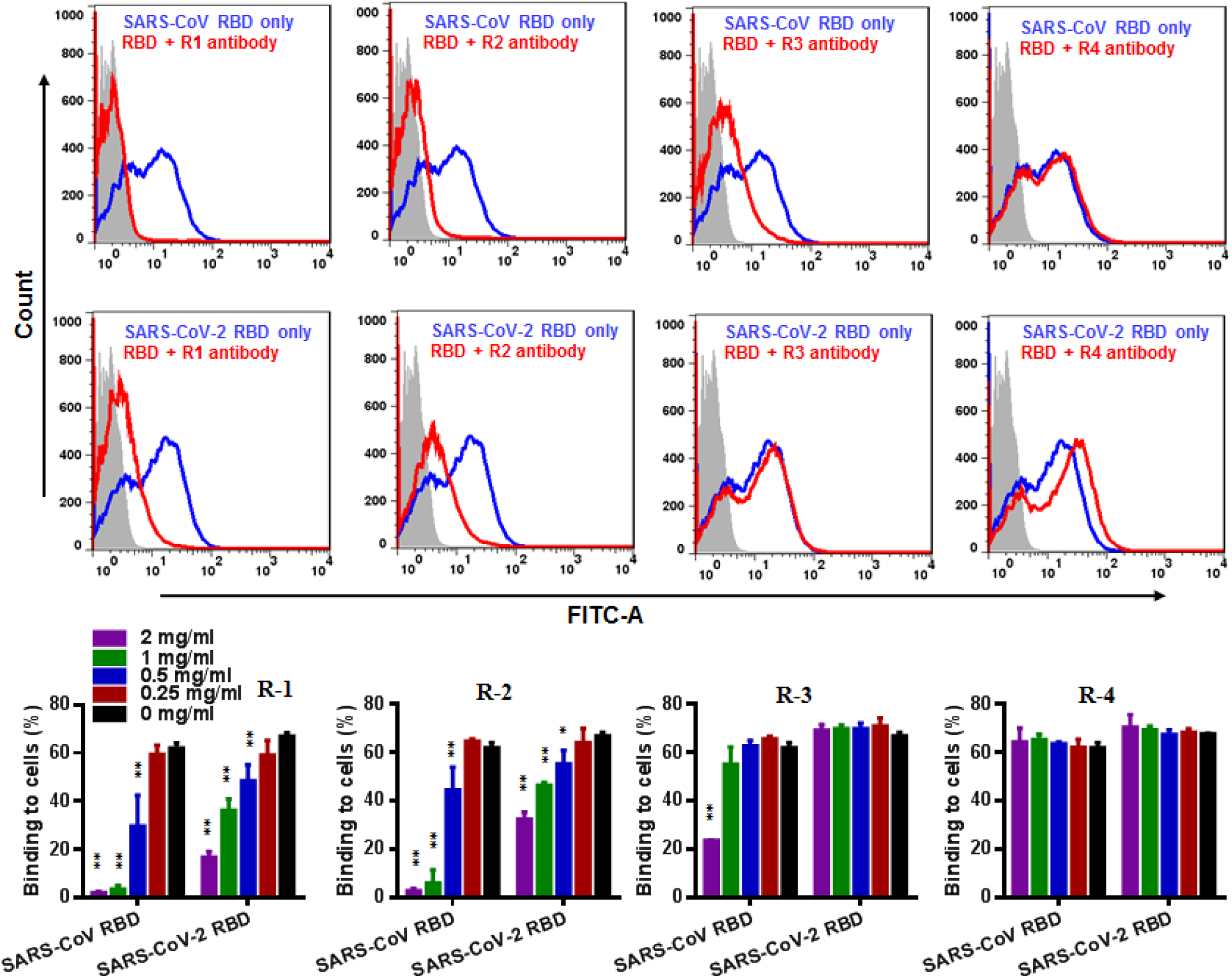
Inhibition of purified rabbit anti-RBD antibodies on the binding of RBD to 293T/ACE2 cells. **(A)** Blocking activity of rabbit anti-RBD antibodies on the binding of SARS-CoV RBD (upper panel) or SARS-CoV-2 RBD (lower panel) to 293T/ACE2 cells was determined by flow cytometry. **(B)** Purified rabbit anti-RBD antibodies inhibited the RBD-ACE2 binding does-dependently. The experiments repeated three times, and data are shown as means with standard deviations. Statistical significance was tested by two-way ANOVA with Dunnett posttest.

## Discussion

To develop effective vaccines and immunotherapeutics against emerging CoVs, the antigenic cross-reactivity between SARS-CoV-2 and SARS-CoV is a key scientific question need be addressed as soon as possible. However, after the SARS-CoV outbreak more than 17 years there are very limited blood samples from SARS-CoV infected patients currently available for such studies. At the moment, Hoffmann *et al*. analyzed three convalescent SARS patient sera and found that both SARS-CoV-2 and SARS-CoV S protein-driven infections were inhibited by diluted sera but the inhibition of SARS-CoV-2 was less efficient (*26*); Qu and coauthors detected one SARS patient serum that was collected at two years after recovery, which showed a serum neutralizing titer of > 1: 80 dilution for SARS-CoV pseudovirus and of 1:40 dilution for SARS-CoV-2 pseudovirus (*27*). While these studies supported the cross-neutralizing activity of the convalescent SARS sera on SARS-CoV-2, a just published study with the plasma from seven SARS-CoV infected patients suggested that cross-reactive antibody binding responses to the SARS-CoV-2 S protein did exist, but cross-neutralizing responses could not be detected (*28*). In this study, we firstly investigated the cross-reactivity and neutralization with a panel of precious immune sera collected from 20 recovered SARS patients. As shown, all the patient sera displayed high titers of antibodies against the S1 and RBD proteins of SARS-CoV and cross-reacted strongly with the S protein of SARS-CoV-2. In comparison, the patient sera had higher reactivity with the S2 subunit of SARS-CoV-2 relative to its S1 subunit and RBD protein, consistent with a higher sequence conservation between the S2 subunits of SARS-CoV-2 and SARS-CoV than that of their S1 subunits and RBDs (*3, 4*). Importantly, each of the patient sera could cross-neutralize SARS-CoV-2 with serum titers ranging from 1:20 to 1:360 dilutions, verifying the cross-reactive neutralizing activity of the SARS patient sera on the S protein of SARS-CoV-2.

Currently, two strategies are being explored for developing vaccines against emerging CoVs. The first one is based on a full-length S protein or its ectodomain, while the second utilizes a minimal but functional RBD protein as vaccine immunogen. Our previous studies revealed that the RBD site contains multiple groups of conformation-dependent neutralizing epitopes: some epitopes are critically involved in RBD binding to the cell receptor ACE2, whereas other epitopes possess neutralizing function but do not interfere with the RBD-ACE2 interaction (*15, 18*). Indeed, most of neutralizing monoclonal antibodies (mAbs) previously developed against SARS-CoV target the RBD epitopes, while a few are directed against the S2 subunit or the S1/S2 cleavage site (*29, 30*). The cross-reactivity of such mAbs with SARS-CoV-2 has been characterized, and it was found that many SARS-CoV-neutralizing mAbs exhibit no cross-neutralizing capacity (*9, 31*). For example, CR3022, a neutralizing antibody isolated from a convalescent SARS patient, cross-reacted with the RBD of SARS-CoV-2 but did not neutralize the virus (*31, 32*). Nonetheless, a new human anti-RBD mAb, 47D11, has just been isolated from transgenic mice immunized with a SARS-CoV S protein, which neutralizes both SARS-CoV-2 and SARS-CoV (*33*). The results of polyclonal antisera from immunized animals are quite inconsistent. For examples, Walls *et al*. reported that plasma from four mice immunized with a SARS-CoV S protein could completely inhibit SARS-CoV pseudovirus and reduced SARS-CoV-2 pseudovirus to ∼10% of control, thus proposing that immunity against one virus of the sarbecovirus subgenus can potentially provide protection against related viruses (*4*); two rabbit sera raised against the S1 subunit of SARS-CoV also reduced SARS-CoV-2-S-driven cell entry, although with lower efficiency as compared to SARS-CoV-S (*26*). Moreover, four mouse antisera against the SARS-CoV RBD cross-reacted efficiently with the SARS-CoV-2 RBD and neutralized SARS-CoV-2, suggesting the potential to develop a SARS-CoV RBD-based vaccine preventing SARS-CoV-2 either (*34*). Differently, it was reported that plasma from mice infected or immunized by SARS-CoV failed to neutralize SARS-CoV-2 infection in Vero E6 cells (*28*), and mouse antisera raised against the SARS-CoV RBD were even unable to bind to the S protein of SARS-CoV-2 (*9*). In our studies, several panels of antisera against the SARS-CoV S and RBD proteins were comprehensively characterized, which provided convincing data to validate the cross-reactivity and cross-neutralization between SARS-CoV and SARS-CoV-2. Meaningfully, this work found that the RBD proteins derived from different SARS-CoV strains can elicit antibodies with unique functionalities: while the RBD from a palm civet SARS-CoV (SZ16) induced potent antibodies capable of blocking the RBD-receptor binding, the antibodies elicited by the RBD derived from a human strain (GD03) had no such effect despite their neutralizing activities. SZ16-RBD shares an overall 74% amino-acid sequence identity with the RBD of SARS-CoV-2, when their internal receptor-binding motifs (RBM) display more dramatic substitutions (∼50% sequence identity); however, SZ16-RBD and GD03-RBD only differ from three amino acids, all locate within the RBM. How these mutations change the antigenicity and immunogenicity of the S protein and RBD immunogens requires more efforts.

Lastly, we would like to discuss three more questions. First, it is intriguing to know whether individuals who recovered from previous SARS-CoV infection can recall the immunity against SARS-CoV-2 infection. For this, an epidemiological investigation on the populations exposed to SARS-CoV-2 would provide valuable insights. Second, whether a universe vaccine can be rationally designed by engineering the S protein RBD sequences. Third, although antibody-dependent infection enhancement (ADE) was not observed during our studies with the human and animal serum antibodies, this effect should be carefully addressed in vaccine development.

## Author contributions

Conceptualization, Y.H., and T.L.; Formal analysis, Y.Z., D.Y., Y.H.; Investigation, Y.Z., D.Y., Y.H., H.C., L.R.; Resources, H.C., L.R., J.W., T.L., Y.H.; Writing-Original Draft, Y.H.; Writing-Review & Editing, all authors; Funding acquisition, Y.H. and T.L..

## Data and materials availability

All data is available in the main text or the supplementary materials.

## Supplementary Materials

## Materials and Methods

### Recombinant S proteins

Two RBD-Fc fusion proteins, which contain the RBD sequence of Himalayan palm civet SARS-CoV strain SZ16 (GenBank: AY304488.1) or the RBD sequence of human SARS-CoV strain GD03T0013 (GenBank: AY525636.1, denoted GD03) linked to the Fc domain of human IgG1, were expressed in transfected 293T cells and purified with protein A-Sepharose 4 Fast Flow in our laboratory as previously described (*15*). A full-length S protein of SARS-CoV Urbani (GenBank: AY278741) was expressed in expressSF^+^ insect cells with recombinant baculovirus D3252 by the Protein Sciences Corporation (Bridgeport, CT, USA) (*16*). A panel of recombinant proteins with a C-terminal polyhistidine (His) tag, including S1 and RBD of SARS-CoV (GenBank: AAX16192.1) and S ectodomain (S-ecto), S1, RBD, and S2 of SARS-CoV-2 (GenBank: YP_009724390.1), were purchased from the Sino Biological Company (Beijing, China).

### Serum samples from recovered SARS patients

Twenty SARS patients were enrolled in March 2003 for a follow-up study at the Peking Union Medical College Hospital, Beijing. Serum samples were collected from recovered patients at 3-6 months after discharge, with the patients’ written consent and the approval of the ethics review committee (*23, 24*). The samples were stored in aliquots at −80°C and were heat-inactivated at 56°C before performing experiments.

### Animal immunizations

Multiple immunization protocols were conducted. First, five Balb/c mice (6 weeks old) were subcutaneously (s.c.) immunized with 20 μg of full-length S protein resuspended in phosphate-buffered saline (PBS, pH 7.2) in the presence of MLP-TDM adjuvant or Alum adjuvant (Sigma-Aldrich). Second, eight Balb/c mice (6 weeks old) were s.c. immunized with 20 μg of SZ16-RBD or GD03-RBD fusion proteins plus MLP-TDM adjuvant. The mice were boosted two times with 10 μg of the same antigens plus the MLP-TDM adjuvants at 3-week intervals. Third, four New Zealand White rabbits (12 weeks old) were immunized intradermally with 150 μg of SZ16-RBD or GD03-RBD resuspended in PBS (pH 7.2) in the presence of Freund’s complete adjuvant and boosted two times with 150 μg of the same antigens plus incomplete Freund’s adjuvant at 3-week intervals. Mouse and rabbit antisera were collected and stored at −40°C.

### Enzyme-linked immunosorbent assay (ELISA)

Binding activity of serum antibodies with diverse S protein antigens was detected by ELISA. In brief, 50 ng of a purified recombinant protein (SARS-CoV S1 or RBD and SARS-CoV-2 S-ecto, S1, RBD, or S2) were coated into a 96-well ELISA plate overnight at 4°C. Wells were blocked with 5% bovine serum albumin (BSA) in PBS for 1 hour at 37°C, followed by incubation with 1:100 diluted antisera or serially diluted purified rabbit antibodies for 1 hour at 37°C. A diluted horseradish peroxidase (HRP)-conjugated goat anti-human, mouse or rabbit IgG antibody was added for 1 hour at room temperature. Wells were washed five times between each step with 0.1% Tween-20 in PBS. Wells were developed using 3,3,5,5-tetramethylbenzidine (TMB) and read at 450 nm after terminated with 2M H_2_SO_4_.

### Neutralization assay

Neutralizing activity of serum antibodies was measured by pseudovirus-based single cycle infection assay. The pseudovirus particles were prepared by co-transfecting HEK293T cells with a backbone plasmid (pNL4-3.luc.RE) that encodes an Env-defective, luciferase reporter-expressing HIV-1 genome and a plasmid expressing the S protein of SARS-CoV-2 (IPBCAMS-WH-01; GenBank: QHU36824.1) or SARS-CoV (GD03T0013) or the G protein of vesicular stomatitis virus (VSV). Cell culture supernatants containing virions were harvested 48 h post-transfection, filtrated and stored at −80°C. To measure the neutralizing activity of serum antibodies, a pseudovirus was mixed with an equal volume of serially diluted sera or purified antibodies and incubated at 37°C for 30 min. The mixture was then added to 293T/ACE2 cells at a density of 10^4^ cells/100 μl per plate well. After cultured at 37°C for 48 h, the cells were harvested and lysed in reporter lysis buffer, and luciferase activity (relative luminescence unit, RLU) was measured using luciferase assay reagents and a luminescence counter (Promega, Madison, WI). Percent inhibition of serum antibodies compared to the level of the virus control subtracted from that of the cell control was calculated. The highest dilution of the serum sample that reduced infection by 50% or more was considered to be positive.

### Flow cytometry assay

Blocking activity of purified rabbit anti-RBD antibodies on the binding of RBD protein with a His tag to 293T/ACE2 cells was detected by flow cytometry assay. Briefly, 2 μg/ml of SARS-CoV-2 RBD protein or 10 μg/ml of SARS-CoV RBD protein were added to 4 × 10^5^ of cells and incubated for 30 min at room temperature. After washed with PBS two times, cells were incubated with a 1:500 dilution of Alexa Fluor^®^ 488-labeled rabbit anti-His tag antibody (Cell Signaling Technology, Danvers, MA) for 30 min at room temperature. After two washes, cells were resuspended in PBS and analyzed by FACSCantoII instrument (Becton Dickinson, Mountain View, CA).

### Statistical analysis

Statistical analyses were carried out using GraphPad Prism 7 Software. One-way or two-way analysis of variance (ANOVA) with Dunnett posttest was used to test for statistical significance. Only p values of 0.05 or lower were considered statistically significant (p>0.05 [ns, not significant], p ≤ 0.05 [*], p ≤ 0.01 [**], p ≤ 0.001 [***]).

---

**Figure S1.**
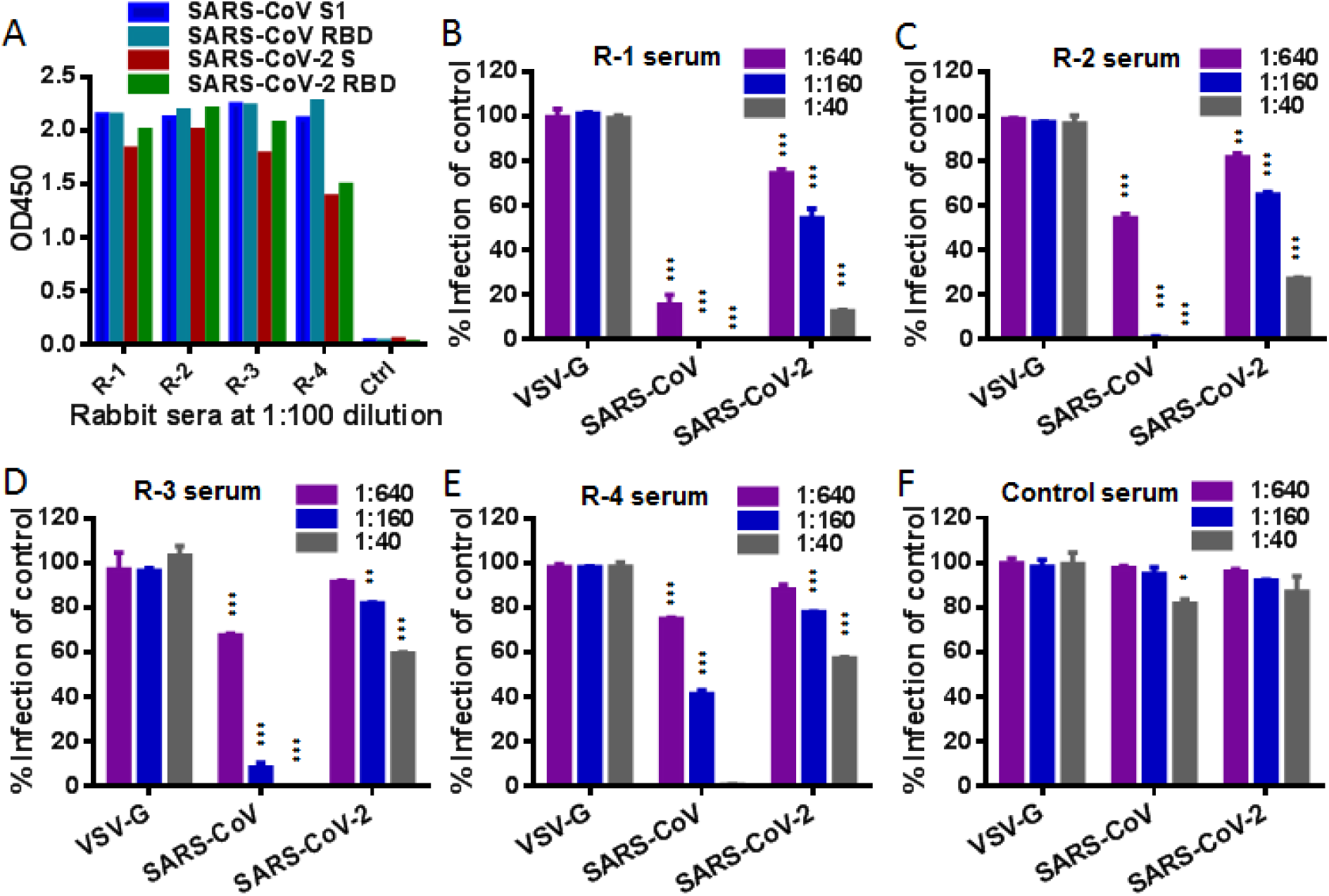
Cross-reactive and neutralizing activities of antisera from rabbits immunized with the RBD proteins of SARS-CoV. **(A)** Binding activity of rabbit antisera at a 1:100 dilution to SARS-CoV (S1 and RBD) and SARS-CoV-2 (S protein and RBD) antigens was determined by ELISA. A healthy rabbit serum was tested as control. **(B)** Neutralizing activity of rabbit antisera or control serum at indicated dilutions on SARS-CoV, SARS-CoV-2, and VSV-G pseudoviruses was determined by a single-cycle infection assay. The experiments were done in triplicates and repeated three times, and data are shown as means with standard deviations. Statistical significance was tested by two-way ANOVA with Dunnett posttest.

**Figure S2.**
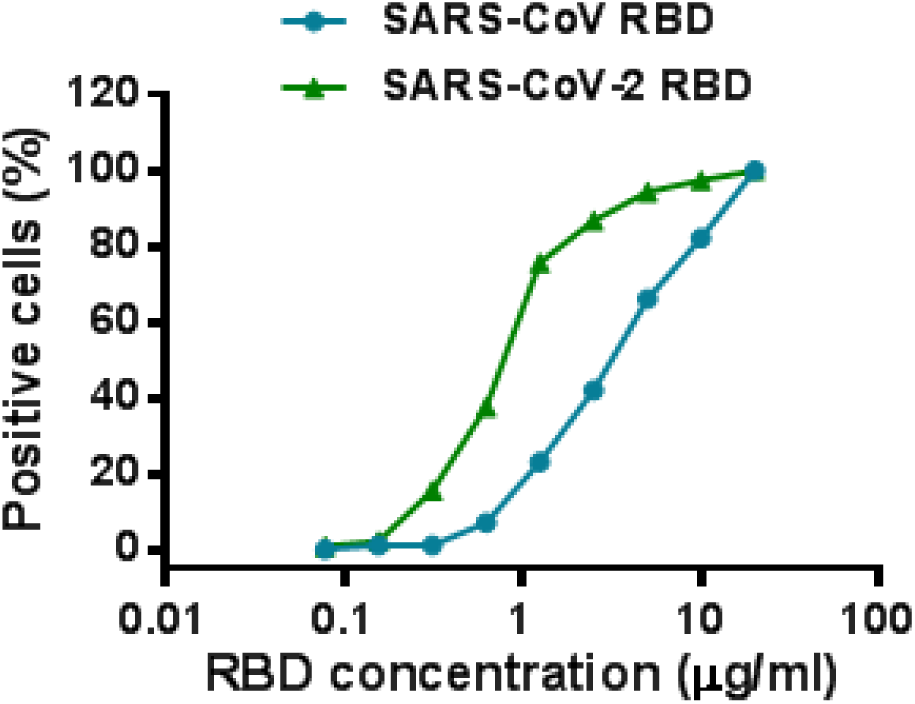
Binding activity of RBD proteins to 293T/ACE2 cells determined by flow cytometry. The assay was repeated two times and obtained consistent results, and representative data are shown.

**Figure S3.**
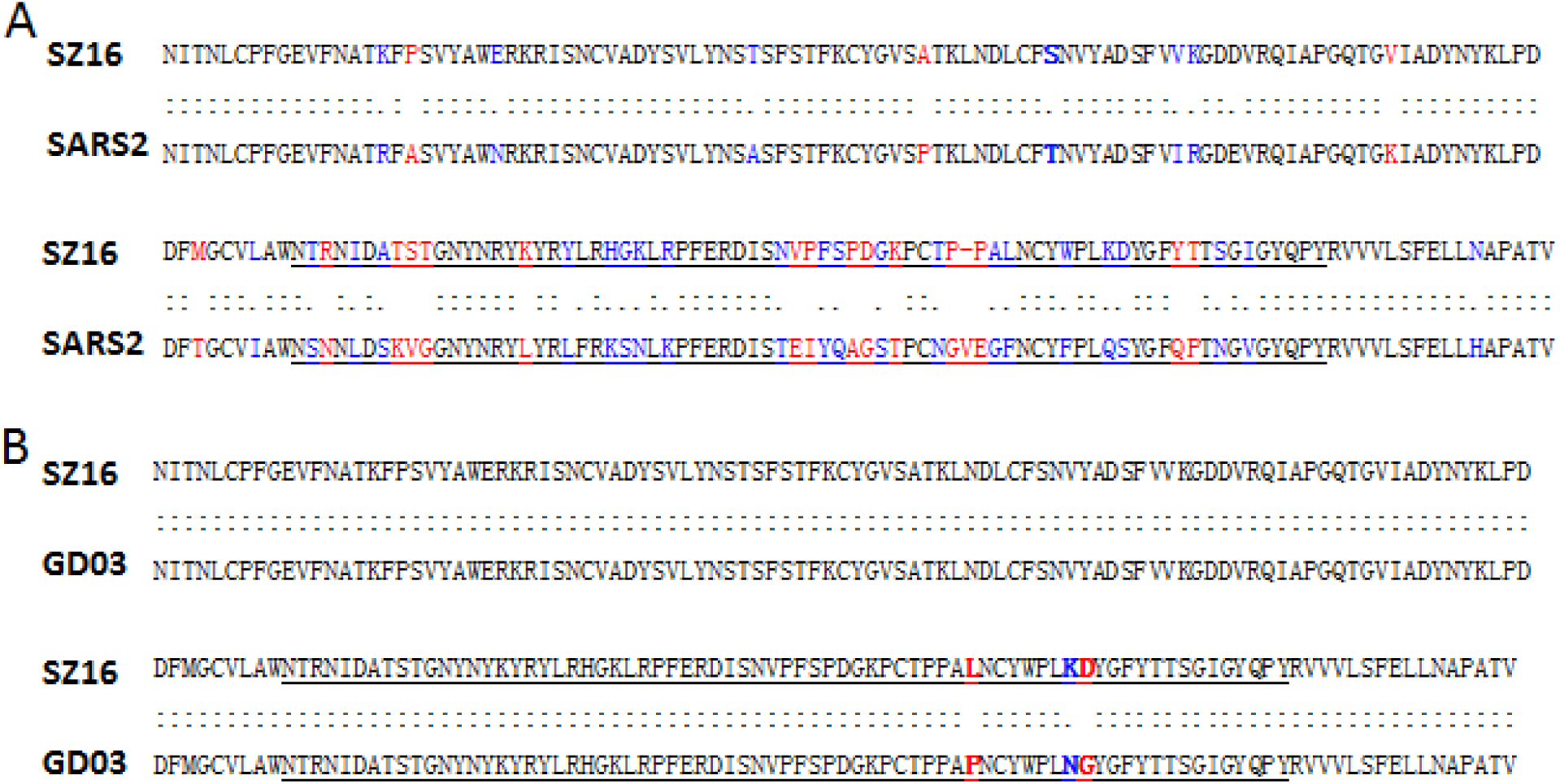
Sequence comparison between the RBDs of SARS-CoV and SARS-CoV-2. **(A)** RBD comparison of the palm civet SARS-CoV strain SZ16 and the human SARS-CoV-2 strain IPBCAMS-WH-01 (designated SARS2). **(B)** RBD comparison of the palm civet SARS-CoV strain SZ16 and the human SARS-CoV strain GD03T0013. Conservative and non-conservative mutations are marked in blue and red, respectively.

